# ChromaFactor: deconvolution of single-molecule chromatin organization with non-negative matrix factorization

**DOI:** 10.1101/2023.11.22.568268

**Authors:** Laura M. Gunsalus, Michael J. Keiser, Katherine S. Pollard

## Abstract

The investigation of chromatin organization in single cells holds great promise for identifying causal relationships between genome structure and function. However, analysis of single-molecule data is hampered by extreme yet inherent heterogeneity, making it challenging to determine the contributions of individual chromatin fibers to bulk trends. To address this challenge, we propose ChromaFactor, a novel computational approach based on non-negative matrix factorization that deconvolves single-molecule chromatin organization datasets into their most salient primary components. ChromaFactor provides the ability to identify trends accounting for the maximum variance in the dataset while simultaneously describing the contribution of individual molecules to each component. Applying our approach to two single-molecule imaging datasets across different genomic scales, we find that these primary components demonstrate significant correlation with key functional phenotypes, including active transcription, enhancer-promoter distance, and genomic compartment. ChromaFactor offers a robust tool for understanding the complex interplay between chromatin structure and function on individual DNA molecules, pinpointing which subpopulations drive functional changes and fostering new insights into cellular heterogeneity and its implications for bulk genomic phenomena.

## Introduction

Chromatin is intrinsically dynamic, and its behavior across time restricts and permits the precise regulatory landscape controlling gene expression ^1,2^. Recent single-cell technologies such as single-cell Hi-C ^3,4^ and chromatin microscopy techniques ^5–9^ now offer unique insight into genome folding, allowing us to directly observe chromatin folding as well as functional readouts in individual cells to disentangle their mechanistic relationship.

Linking chromatin conformation to function in single cells presents several key challenges: 1) Single cell data is extremely sparse ^10^. Current single-cell technology often yields incomplete information, such as missing values or misallocated genomic coordinates. 2) Single-cell measurements capture snapshots, whereas chromatin function may result from a dynamic behavior as it moves across time. Phenomena observed in bulk experiments, such as Hi-C, may be artifacts of averaging across cell populations and patterns seen in bulk may not exist in single cells ^11^. 3) Capturing chromatin folding and phenotypic measurements like nascent transcription in the same cells has only recently become possible, but temporal offsets between folding and function could introduce uncertainty. 4) The bulk trends we observe in aggregate may be driven by a small fraction of cells, and identifying this subset amidst heterogeneous single-cell chromatin measurements presents a complex challenge. Connecting chromatin behavior to function in individual cells remains intractable given these technical barriers.

Several computational methods have been developed in response to emerging single-cell imaging and high-throughput sequencing techniques to measure chromatin conformation. Topic modeling^12^, random-walk methods^13^ and recent deep learning approaches^14,15^ effectively cluster cells into subpopulations. Recent methods also offer rich annotations in single cells, including A/B compartments ^14,16^, subcompartments^17^, topologically associating domains (TADs)^14,16^, and chromatin loops^18^. Rajpurkar et al. first applied a convolutional neural network to directly predict nascent transcription from chromatin folding^19^ and Zhan et al.^20^ propose an effective deep-learning-based dimensionality reduction method to cluster conformations. However, these works did not yet connect the behavior of individual cells to populations of similar conformations that are transcriptionally on or off. We build on these works to relate the behavior of individual cells to bulk trends as well as mechanistically link chromatin behavior to transcription.

We introduce ChromaFactor, a non-negative matrix factorization (NMF) technique to decompose single-cell datasets into interpretable components and identify key subpopulations driving cellular phenotypes. Non-negative matrix factorization (NMF) offers an ideal application in the analysis of such complex data due to its inherent capacity to reduce high-dimensional data into a lower-dimensional, interpretable format^21^. NMF has a robust legacy in genomics as it allows for the deconvolution of composite signals into a set of additive components and can therefore discern patterns and structures in noisy, large-scale data. Notably, it has been used on bulk Hi-C data for TAD calling^22^ and has found applications in other emergent single-cell modalities, including single-cell RNA-Seq^23^ and spatial transcriptomics datasets^24^. By applying NMF to single-cell genome folding datasets, we can identify significant components or *‘templates’* that account for the majority of cellular variation. Linking these templates to matched functional readouts describes how differences in cell populations correspond to differences in phenotypes. ChromaFactor deconvolves single-cell chromatin organization datasets into their most meaningful primary components, providing new insights into the interplay between chromatin structure and function. Here, we apply ChromaFactor to two single-cell imaging datasets and link templates to nascent transcription. This tool may also be applied to any set of ordered matrices in single cells.

## Results

### NMF to decompose single-cell 3D genome conformation datasets

We were motivated to develop ChromaFactor by the disconnect between meaningful signal observed bulk cell populations and the extreme heterogeneity of single-molecule examples. One such dataset, Mateo et al. 2019^5^, profiles local chromatin conformation at the bithorax complex (BX-C) in *Drosophila* embryos and additionally includes matched nascent transcription in the same cells (*n* = 19,103). To discover how cell populations vary, we often take the difference between the average contact maps under two conditions. We observed a pronounced boundary in cells within the 30 kb region actively transcribing the Abd-A gene, as compared to non-transcribing cells (**Fig. 1a**). However, these patterns are nearly impossible to discern in individual cells (**Fig. 1b**). Given the heterogeneity and dynamic behavior of chromatin in single cells, identifying which cells contribute to overall trends is complex. Are these contact patterns visible at the single-cell level, or are they composite effects resulting from population-wide averaging? To bridge the gap between single-cells and bulk averages and identify which cells contribute most, we propose applying NMF to single-molecule chromatin conformation datasets with ChromaFactor.

**Figure 1:**
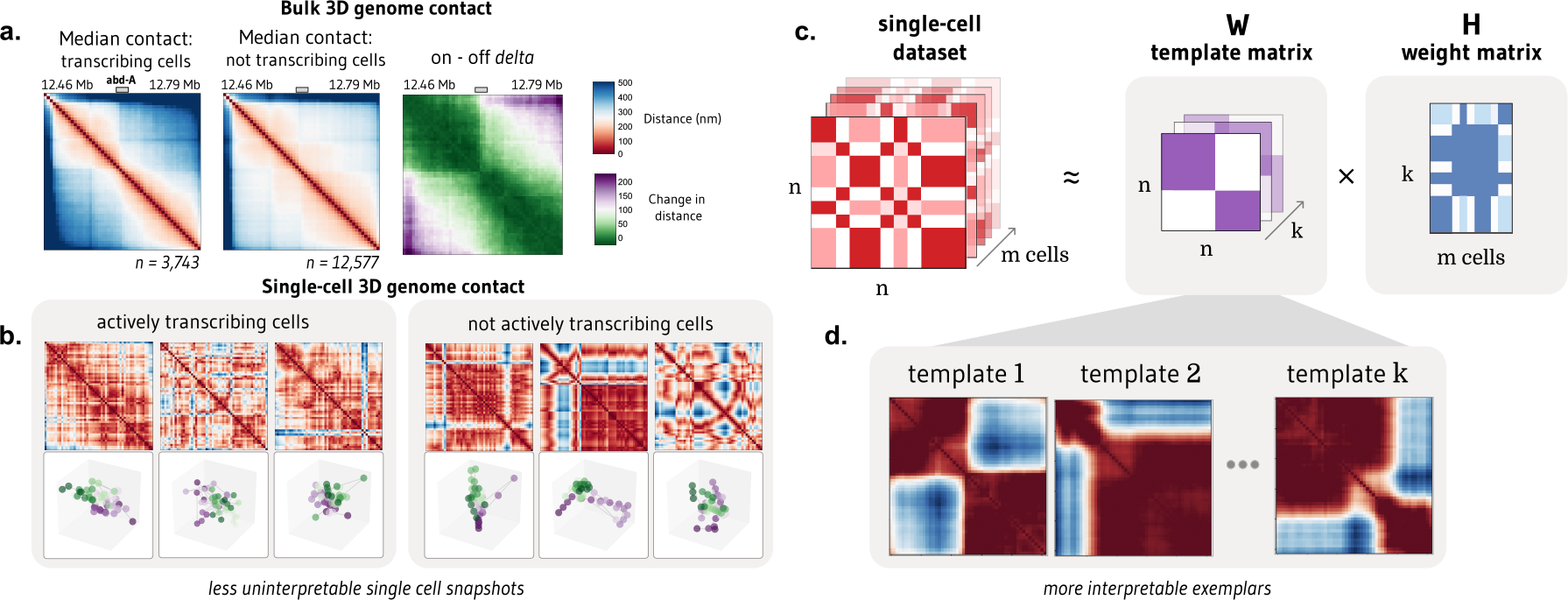
NMF provides interpretable decomposition of single-molecule chromatin conformation datasets. **a**. Matrices representing the median all-by-all Euclidean difference (nm) between genomic loci in single molecules at a 30 kb locus in *Drosophila melanogaster* actively transcribing (left) and not transcribing (middle) the Abd-A gene from Mateo et al. (*n* = 16,320 molecules). The rightmost panel shows the difference in distance matrices, indicating two domains with elevated local interactions and reduced distal interactions in populations transcribing Abd-A. **b**. Bulk trends in contact change are challenging to observe in single cells actively transcribing (left) and not transcribing (right) Abd-A. **c**. Non-negative matrix factorization (NMF) decomposes a dataset of single-cell distance matrices into a *template* matrix with interpretable chromatin domain boundaries and a *contribution* matrix describing the weight of each template to each cell. **d**. Three templates produced when NMF is applied to distance matrices at the Abd-A locus.

In this approach, NMF decomposes a non-negative distance matrix into two lower-rank non-negative matrices, such that their product approximates the original matrix. ChromaFactor decomposes an *n* by *n* by *m* count matrix, where *n* is the number of genomic loci profiled and *m* is the number of cells, into an *n* by *n* by *k* component matrix W, where *k* is a specified number of components, and a *k* by *m* weight matrix, H (**Fig. 1c**). The matrix W represents the basis vectors, which we call *templates*, as they resemble patterns observed across the cell population. The method accepts both distance matrices from single-molecule imaging experiments and contact matrices from single-cell Hi-C. Here, the number of components (*k*) was selected to balance template interpretability and reconstruction error. The matrix H represents the weight matrix, signifying contributions of cells to components such that the data for each molecule is approximated as the weighted average of the components plus noise. To estimate W and H, matrices are randomly initialized and updated to minimize the reconstruction error between their product and the single-cell dataset.

### Relating single cells to bulk trends with ChromaFactor

When ChromaFactor is applied to the Mateo et al. dataset with twenty components, we find that several templates resemble chromatin boundaries (**Methods, Fig. 1d, Supp. Fig. 1, Supp. Fig. 2**). To visualize the relationships within the single-cell dataset, we apply UMAP on the weight matrix, H (**Fig. 2a**), and label cells by the component with the largest weight. Investigating individual examples, we find that single cells can resemble these template patterns. To illustrate, we show the 3D coordinates and distance maps of three cells, along with their component contributions from weight matrix H (**Fig. 2b-d**). Since every cell is an additive combination of the templates, we can multiply each cell’s weights by the component matrix to reconstruct single-cell examples, which are less noisy than the original cell measurements (**Fig. 2e**). Notably, these cells closely resemble the component with the highest contribution, indicating that templates can be representative of cell subpopulations (**Fig. 2f**). Indeed, considering all cells in the same group, we find that median contact resembles the closest template (**Fig. 2g**)

**Figure 2:**
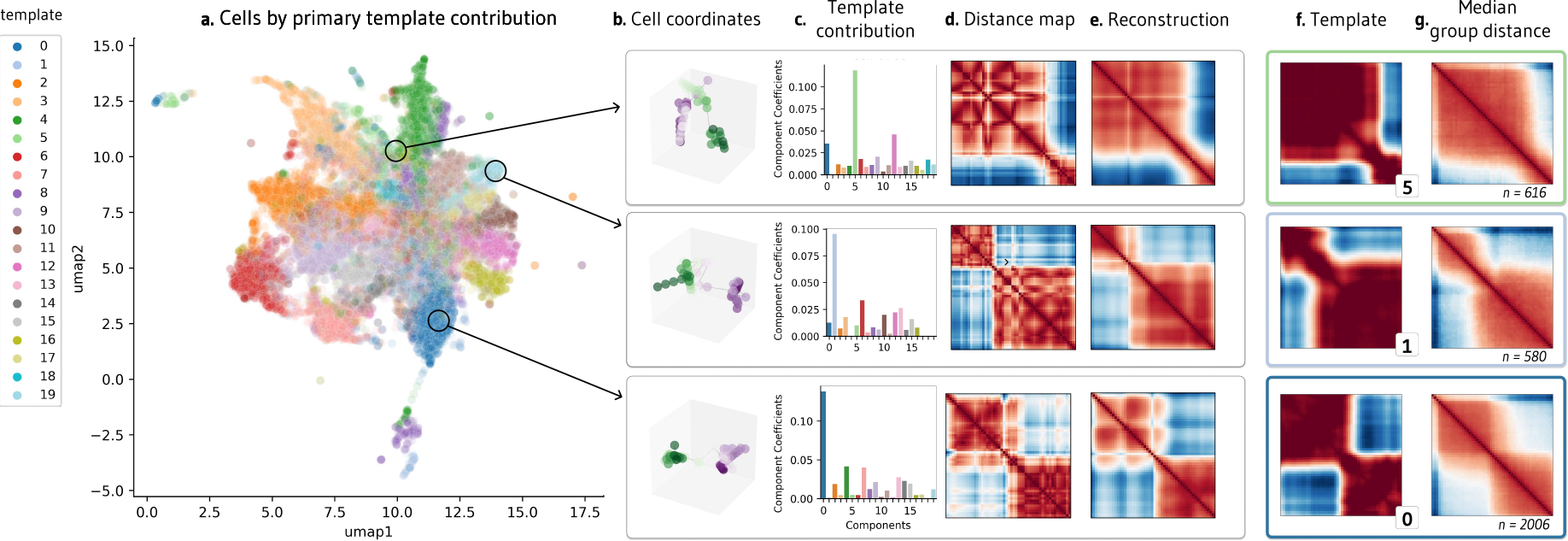
Visualization of NMF outputs and their relationship to single-cell behavior. **a**. UMAP visualization of contribution matrix, colored by the template with the predominant contribution in each cell. **b**. Depiction of cell coordinates from selected individual cells. **c**. Component contributions for each cell, emphasizing high weight for templates 5, 1, and 0. **d**. Distance matrices corresponding to each cell. **e**. Denoised reconstructions of the distance matrices, created by multiplying the template contribution of a cell shown in (c) by the template matrix. **f**. NMF templates 5, 1, and 0, which had the highest weight contributions for the three individual cells in (c). **g**. Median contact distance across all cells in the dataset with the highest weight contributions to templates 5, 1, and 0, respectively.

### Templates are significantly correlated with active transcription

Templates may capture subsets of cells and cellular patterns. Which components, if any, correspond to biological phenotypes? To investigate if templates are correlated with downstream biological function, we train a random forest model to predict nascent transcription of nearby genes from the weight matrix H alone (**Methods, Fig. 3a**). In the Mateo et al. dataset, three genes were profiled in the same cells that were imaged, producing matched chromatin organization and transcription data^5^. Predictive performance would indicate that the components capture salient information about transcriptional state and may serve as a proxy for the raw input distance matrices themselves.

**Figure 3:**
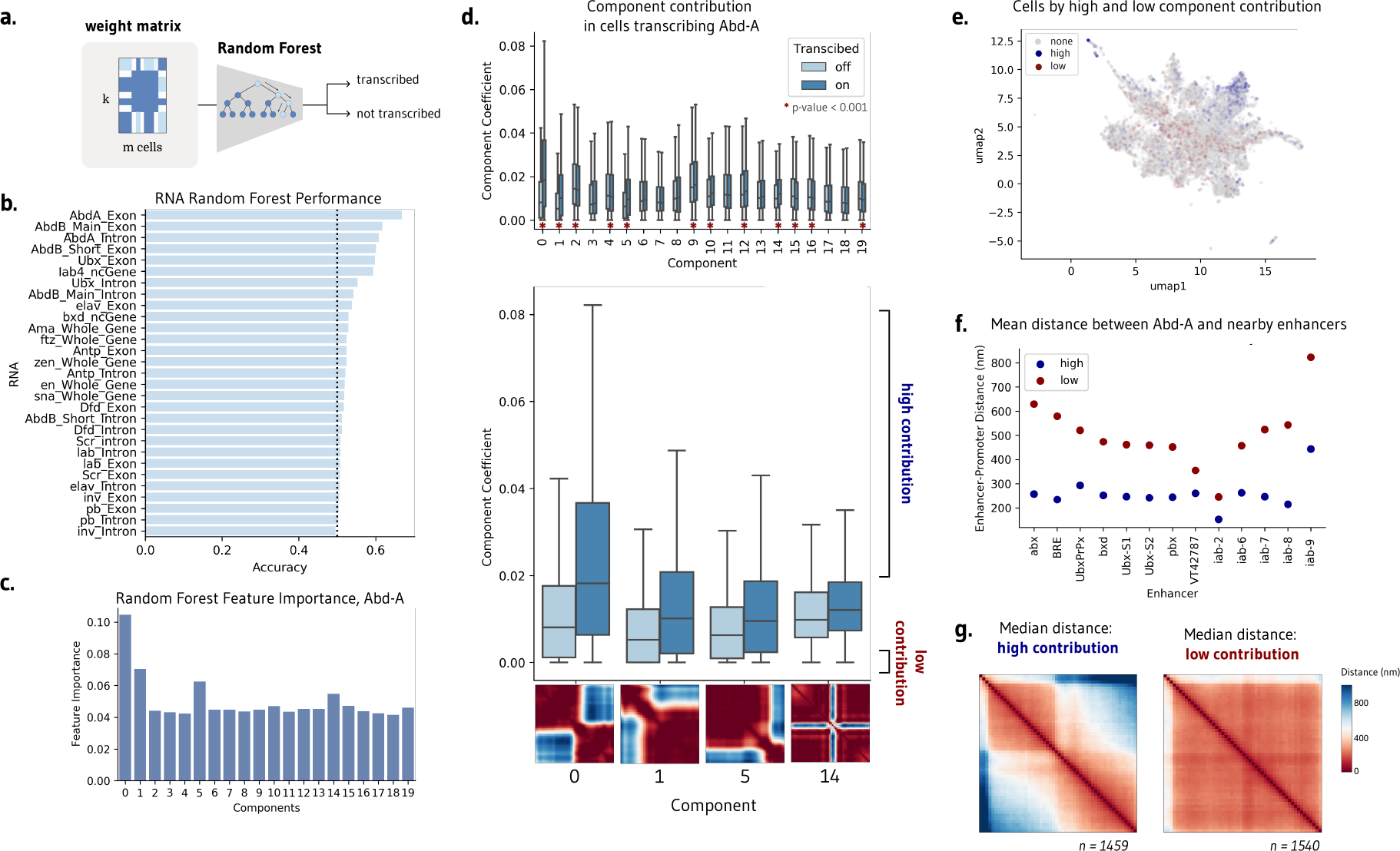
NMF templates are significantly correlated with transcription. **a**. Application of random forest models to predict cell transcription from the contribution matrix alone. **b**. A random forest can modestly predict transcription in abd-A, Abd-B, and Ubx, demonstrating that the components capture salient information for transcription. **c**. Random forest feature importance highlights templates 0, 1, and 5 as most important for predicting transcription. **d**. Several components, including 0, 1, 5, and 14, have significantly different component contribution weights in transcribing and not transcribing cells. **e**. UMAP visualization of component contribution matrix, colored by cells with a high contribution of components 0, 1, 5, and 14 (blue) and a low contribution of these components (red). **f**. Mean distance between abd-A and nearby enhancers at the same locus across the subset of cells with high and low component contributions. **g**. Median contact of cells with high and low component contributions, encompassing the subset of cells which may be responsible for changes in contact observed in bulk.

Different random forest models were separately trained to predict transcription in the 17 measured gene isoforms. We find that the weight matrix can modestly predict transcription across several genes, including Abd-a, Ubx, and Abd-b on balanced datasets of transcribing and non-transcribing cells (**Fig. 3b**). Indeed, the performance of the random forest parallels the performance of a random forest trained directly on the distance matrices, achieving an accuracy of 67.4% and 65.3%, respectively. Examining the feature importance of the Abd-a trained model, we find that components 0, 1, 5, and 14 are particularly influential for the model’s prediction (**Fig. 3c**).

We can alternatively address which components are preferentially upweighted by transcribed cells by evaluating component weights separately for transcribing and non-transcribing cells. Twelve of the twenty components have significantly different weights between Abd-A transcribing and non-transcribing cells (two-sided Mann Whitney U, p-value < 0.001, **Fig. 3d**). This effect is most extreme in components 0, 1, 5, and 14, the same components identified as salient by the random forest models. Visually, these components show a separation of chromatin into two distinct compartments across three separate points across the locus (components 0, 1, 5), as well as a sharp decrease in contact at the center of the locus (component 14). Significant templates suggest how contact differs upon active transcription to favor stricter subcompartmentalization within the genomic locus.

### Subpopulations of transcribing cells drive contact patterns observed in aggregate

Our aim is to understand not just how contact differs, but which subpopulations of cells drive the changes we observe in bulk (**Fig. 1a**). We consider transcribing cells with the top 50% of weights in components 0, 1, 5, and 14, which we call ‘high contribution’ cells, and contrast them with non-transcribing cells in the bottom 50% of component contributions (‘low contribution’ cells). These cells make up only 12.7% of the total cell population, but their component weights are the most predictive of transcriptional state. These high and low contribution cells occupy different areas of the UMAP plot seen previously– low contribution cells are more likely to be mixes of components and high contribution cells are more likely to favor one component, suggesting that biologically consequential cells resemble templates (**Fig. 3e**).

These subpopulations of cells differ not just in chromatin conformation and transcriptional state, but also in their local behavior of regulatory elements. The distance between Abd-A and all proximate enhancers at the same locus is notably smaller in high contribution cells as compared to low contribution cells (**Fig. 3f**). Enhancers are closer to the gene promoter in the subset of active cells identified by ChromaFactor. Moreover, examining the median contact of high and low contribution cells, we observe contact patterns far more pronounced than those observed in bulk (**Fig. 3g**). Cells with high component weights possess stronger boundary separation as well as a stripe of contact centered at the location of Abd-A, which is absent in low-contribution, transcriptionally-off cells. In this case, the cell population identified by ChromaFactor exhibits a more potent and unified profile of compartmentalized chromatin driving smaller enhancer-promoter distances when compared with all transcribing cells.

In sum, template analysis at this locus paints a holistic portrait of higher genomic sequestration between loci, reducing the distance between enhancers and promoters, thereby increasing the likelihood of transcription. This trend, although suggested at the level of the bulk population, is strongly driven by a small subpopulation of single cells. The remaining population is extremely heterogeneous across transcribing and non-transcribing cells such that their contact effectively cancels out.

### Application of ChromaFactor to holocarboxylase synthetase (HLCS) locus highlights local and compartment-level chromatin shifts upon transcription

To demonstrate the efficacy of ChromaFactor across genomic scales, we next apply NMF to a 10 Mb locus in human IMR90 cells with 40 kb resolution^6^. We profile a population of 7,590 cells derived from genome-wide profiling of chromatin conformation and nascent transcription with microscopy. The median genomic distance between cells actively transcribing and not transcribing the HLCS gene reveals no visually discernible change in contact (**Fig. 4a**). However, after subtracting one contact matrix from the other to examine the difference in contact between populations, we observe a weakened boundary directly upstream of the HLCS locus.

**Figure 4:**
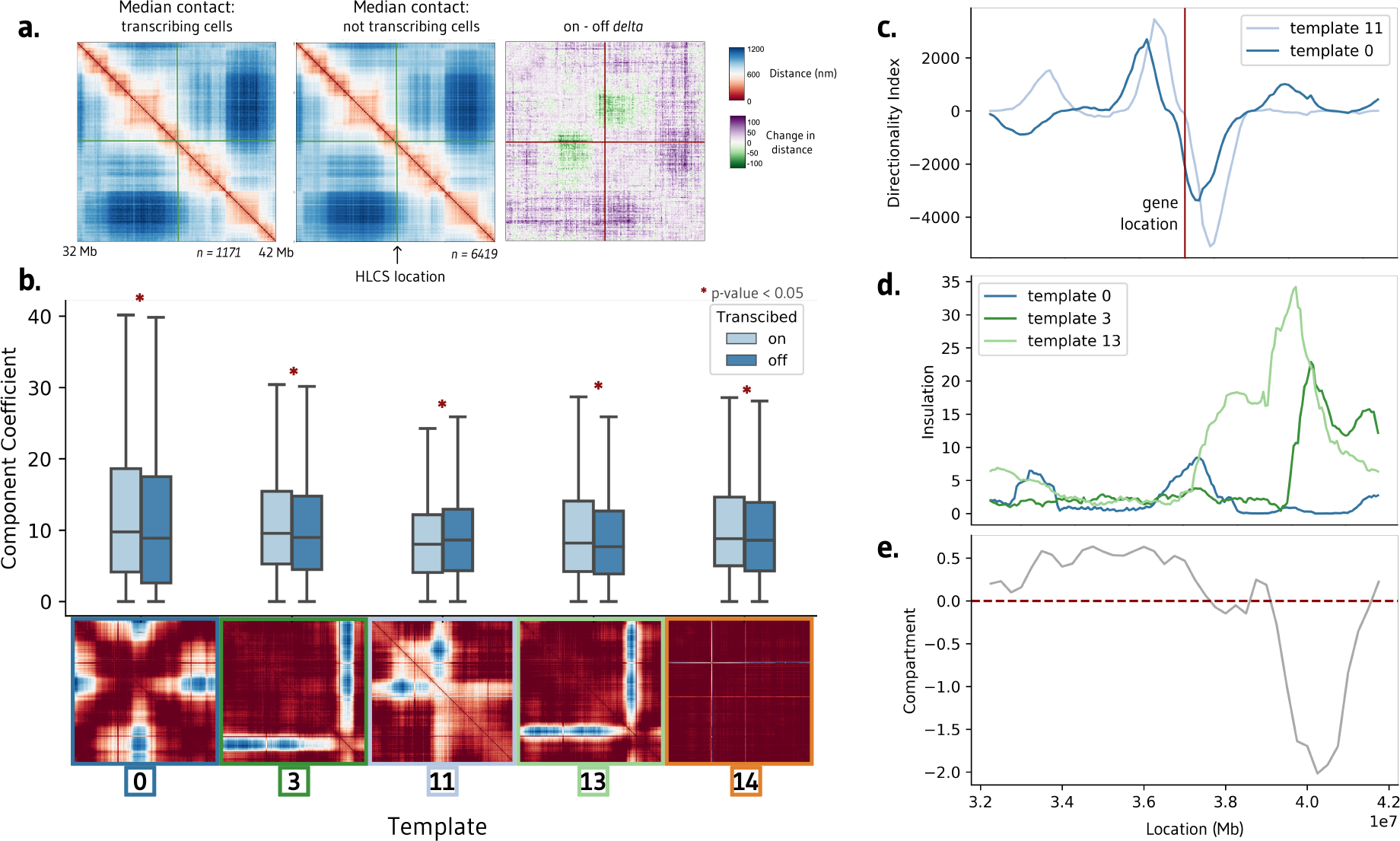
Interpretable templates at the HCLS locus in IMR-90 cells. **a**. Average chromatin contacts in cells actively transcribing (left) and non-transcribing (middle) the HCLS gene within the surrounding 10 Mb region. The right panel highlights the contrast in contact patterns, emphasizing a stronger boundary in actively transcribing cells. **b**. Templates generated using Non-Negative Matrix Factorization (NMF) on this cell population. Among the 20 components, five exhibit significant differences between cells transcribing and not transcribing the HCLS gene. **c**. Directionality index of templates 0 and 11 correspond with the location of the transcribed HLCS gene. **d-e**. Insulation scores of templates 3 and 4 align with a shift in compartment in IMR-90 cells.

We decompose the single-cell imaging dataset into twenty components with ChromaFactor and identify components with significantly different weights in transcribing cells (**Fig. 4b, Supp. Fig. 3**). Of the twenty components, five are significantly different in transcribed cells (two-sided Mann-Whitney-U, p-value < 0.05), exhibiting a diverse range in contact differences across templates. Component weights are elevated in transcribing cells in all components but component 11, where a decrease in contact is observed at the precise location of the boundary loss observed in bulk. Curiously, component 14 highlights a sharp change in contact at two particular genomic loci, one of which is the location of the HLCS gene itself. The method has no knowledge of transcriptional state nor gene location, indicating that change in contact at this locus may nonetheless contribute to a significant amount of variation across the cell population.

We find that two significant components, 0 and 11, correspond to a steep shift in the directionality index, a measure of contact frequency bias towards either upstream sequence or downstream sequence, at the location of the HLCS locus (**Fig. 4c**). Remaining components 3 and 13 display a sharp increase in insulation at the location of a compartment switch from A to B in IMR90 cells (**Fig. 4d-e**). In sum, template analysis at this locus suggests that the change of transcription correlates with a reorganization of chromatin directly at the site of active transcription, at boundary shifts directly upstream of the locus, and within broader compartments downstream of the locus.

## Discussion

This study has demonstrated the utility of ChromaFactor, a novel application leveraging Non-negative Matrix Factorization (NMF) for dissecting single-cell chromatin conformation datasets. Our method uncovers nuanced layers of genome conformation dynamics and their correlation with transcriptional states, which would otherwise be obscured in bulk analyses. Correlations between template patterns and active transcription suggest that these templates are not merely reflections of cellular heterogeneity, but could be mechanistically linked to transcriptional regulation. Our application of ChromaFactor to the Mateo et al. and Su et al. datasets leads us to two intriguing biological interpretations: 1) bulk behavior can sometimes be observed in individual cells, and 2) only a small minority of cells in the population drive population-wide signal. This reasoning is not possible at the bulk level, where trends may be artifacts of averaging, nor at the single-cell level, where it is unknown which snapshots capture relevant signal.

Looking ahead, the application of ChromaFactor to a wider array of cell types, genomic phenomena, and sc-HiC will help us to understand if these conclusions hold across biological contexts. ChromaFactor’s ability to isolate functional portions of single-cell chromatin datasets could clarify the structural dynamics underpinning cell-type-specific gene regulation and the development of cellular heterogeneity. Additionally, integrating ChromaFactor with multi-omics approaches, such as single-cell RNA-Seq, ATAC-Seq, or CUT&Tag, could help resolve the interplay between chromatin structure, epigenetic modifications, and gene expression. ChromaFactor’s application to pathological states or CTCF degradation experiments presents another exciting area of exploration.

While the application of ChromaFactor is promising, it is important to note its current limitations. The number of templates, *k*, is manually selected. The approach also relies heavily on the quality of the single-cell chromatin datasets and the resolution at which they are produced. The inherent noise and technical artifacts present in these datasets can influence the deconvolution process and the interpretation of the resulting components. Extremely noisy datasets with consistent dropout locations will produce templates capturing dropout. Further improvements in single-cell chromatin imaging and sequencing technologies will likely enhance the accuracy and interpretability of ChromaFactor’s outputs.

Finally, a deeper investigation of the biological interpretation of components is warranted. While we have shown that these components correlate with transcription and other genomic features, the exact mechanisms through which these templates influence cellular behaviors remain largely unknown. Additional studies are needed to mechanistically link these structural templates with specific functional outputs and to explore their potential role in modulating regulatory response. In sum, this study introduces ChromaFactor as a promising tool for decoding single-cell chromatin conformation data. It provides a more granular view of the dynamic nature of genome architecture and its role in gene regulation, thereby broadening our perspective on the intricate interplay of genome organization and function.

## Code Availability

This package as well as notebooks to reconstruct the figures can be found at the following repository: https://github.com/lgunsalus/ChromaFactor.

## Acknowledgments

We thank Chris Olah for the original idea to apply NMF to single-cell 3D genome datasets and Kangway Chuang for project guidance, including the idea to train a random forest to test component significance. We additionally thank Archit Verma, Amanda Everitt, Katie Gjoni, Shuzhen Kuang, and other members of the Pollard and Keiser labs for helpful discussion and manuscript feedback.

## Funding Sources

This work was supported by the NIH 4D Nucleome Project (grant #U01HL157989 to K.S.P.), grant number 2018-191905 from the Chan Zuckerberg Initiative DAF (M.J.K.), an advised fund of the Silicon Valley Community Foundation (M.J.K.), and a UCSF Achievement Rewards for College Scientists (ARCS) Scholarship (L.M.G.).

## Methods

### Datasets and processing

#### Mateo et al. dataset

The Mateo et al. microscopy dataset contains 3D genomic coordinates and transcriptional activity for single molecules spanning the Drosophila Bithorax complex (BX-C) locus^5^, which can be found at the following repository: https://zenodo.org/records/4741214. We employed the data preprocessing procedure from Rajpurkar et al.^19^ to handle missing values, which can be found at the following repository: https://github.com/aparna-arr/DeepLearningChromatinStructure/tree/master/DataPreprocessing.

Cells with over 80% of coordinates missing were excluded from our analysis. For the remaining cells, missing coordinates were imputed by linear interpolation between adjacent loci using the scipy.interpolate.interp1d function^25^. Maps were normalized by dividing by the maximum distance observed as followed prior to NMF.

#### Su. et al. dataset

The Su et al. dataset^6^ comprises genome-wide chromatin folding data from single molecules in human IMR90 fibroblasts imaged using DNA FISH. Data can be downloaded from the following repository: https://zenodo.org/record/3928890. In particular, we analyze paired coordinate and transcription data in ‘genomic-scale-with transcription and nuclear bodies.tsv’. Custom Python code was written to extract specific genomic regions from the raw dataset, transform coordinate data into distance matrices, and identify cells transcribing the HLCS (ENSG00000159267) gene for downstream analysis, and can be found in the provided repo with an example of processing. Maps were considered with a 40k resolution. Cells with more than 25% missing coordinates within the 10 Mb HLCS genomic locus were excluded. Any remaining missing values were imputed by linear interpolation using numpy.interp^26^.

### Non-negative matrix factorization (NMF) and ChromaFactor application

We applied non-negative matrix factorization (NMF) using the ChannelReducer wrapper in the Lucid NMF library^27^ built on top of the scikit-learn implementation^28^, which unravels the 3D input matrices into 2D vectors suitable for NMF. We set the number of components to k=20 to balance interpretability of templates and reconstruction error (**Supplementary Figure 1**). Default scikit-learn NMF parameters were used: NNDSVD initialization, a coordinate descent solver, Frobenius loss, tolerance of 1e-4, maximum 200 iterations, and an element-wise L2 regularization penalty. Additional code is provided to process both datasets and plot 2D distance matrices and 3D coordinates.

### Random forest

We trained random forest classifiers using RandomForestClassifier in scikit-learn^28^ to predict transcriptional activity from NMF component weights. Models were trained separately for each gene with binary on/off transcription labels. Balanced datasets were created for each gene with equal transcribing and non-transcribing cells. Data was split 70/30 into train and test sets. All other random forest parameters were left as scikit-learn defaults.

### Genomic features annotations

Gene annotations and enhancer locations were derived from the original publications for each dataset. Enhancer and gene locations were provided from Mateo et al^5^. Compartment annotations were used from Rao et al.^29^ (4DNFIHM89EGL). Directionality index and insulation tracks were produced from scoring code provided in Gunsalus et al^30^. UMAP dimensionality reduction for visualization used the scikit-learn implementation with n_neighbors=5 and remaining default parameters.

### Statistical Analysis

Differences in contact patterns and NMF component weights between transcribing and non-transcribing cells were evaluated using the non-parametric two-sided Mann-Whitney U statistical test. P-values less than 0.05 were considered statistically significant.

## Notes

### Competing Interest Statement

The authors have declared no competing interest.

