## Supplemental for "ChromaFactor: deconvolution of single-molecule chromatin organization with non-negative matrix factorization"

### ChromaFactor Supplementary Information

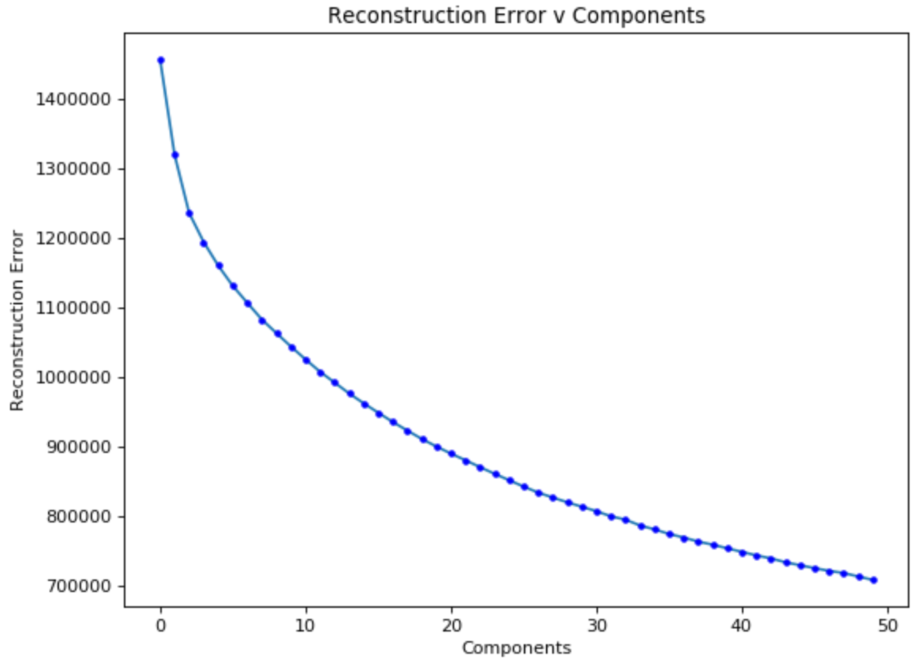


**Supplementary Figure 1: Reconstruction error across number of components, *k*.** Reconstruction error, measured by the difference between the original single-cell chromatin folding dataset and the NMF reconstructed approximation, across different values of *k* components. Adding more components reduces the reconstruction error as NMF can better capture patterns in the data at the expense of interpretability. An elbow is visible around k=15-20 where adding additional components leads to diminishing returns in error reduction. We selected k=20 components for our analysis to balance reconstruction accuracy and interpretability.

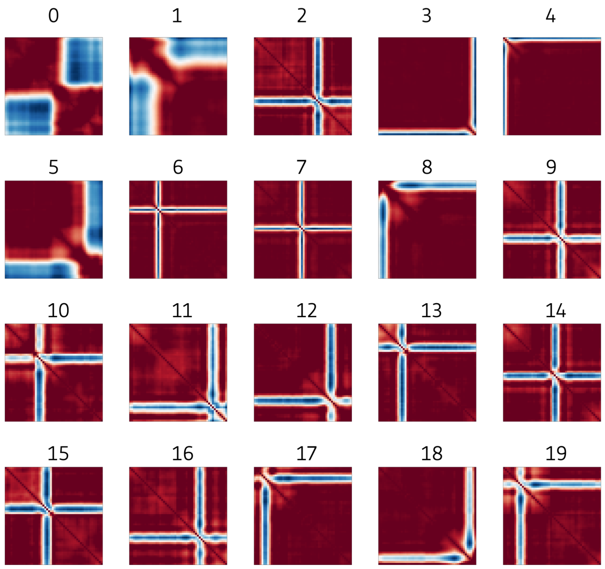


**Supplementary Figure 2: Components at BX-C locus.** All 20 components generated by applying NMF across the cells at this locus from the Mateo et al. dataset.

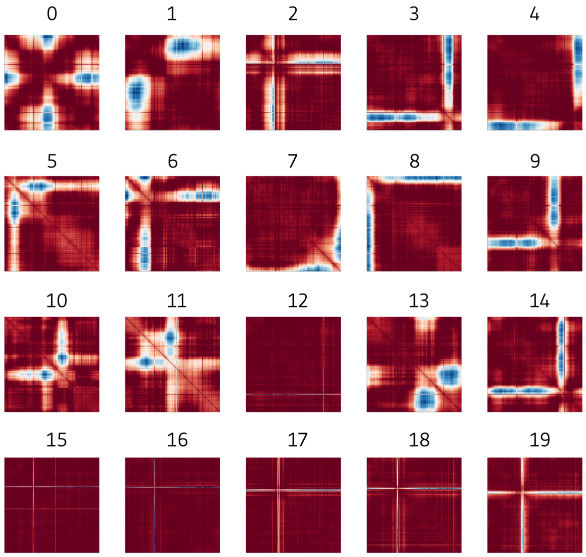


**Supplementary Figure 3: Components at HLCS locus.** All 20 components generated by applying NMF across the cells at this locus from the Su et al. dataset.
